# Modeling the Neurocircuitry Involved in Binge Drinking

**DOI:** 10.64898/2026.09.28.754793

**Authors:** Amy Rude, Nicholas Hayek, Christopher Lapish, Alexey Kuznetsov

## Abstract

Binge drinking is characterized by an initial bout of rapid, high-vigor alcohol intake (front-loading) followed by a prolonged period of lower-rate “maintenance” drinking, yet the circuit mechanisms that generate and regulate these phases remain unclear. Here, we develop a systems-level computational model that links the structure of binge drinking to interactions among medial prefrontal cortex (mPFC), insula, striatum, and ventral tegmental area (VTA). The model consists of cortical Seek (mPFC) and Go (insula) populations that form a mutually excitatory loop, a SetPoint population that integrates striatal activity as an internal estimate of effort/consumption, ventral (nucleus accumbens, NAc) and dorsal (dorsolateral striatum, DLS) striatal populations, and a dopaminergic VTA population providing transient reward-prediction signaling. Using a firing-rate formalism with noisy dynamics, we reproduce the canonical two-phase binge pattern and show that: (i) the PFC–insula loop acts as a bistable module that initiates and sustains front-loading; (ii) striato-cortical feedback to the Setpoint population controls the timing of the transition from front-loading to maintenance without altering its vigor; and (iii) the amplitude of the VTA dopamine transient selectively scales front-loading intensity. Sensitivity analyses identify PFC–insula connectivity, VTA-to-NAc modulation, and striato-cortical feedback as key parameters that bifurcate two-phase drinking into regimes with abolished or persistent front-loading. These findings provide a mechanistic framework for interpreting binge-like alcohol intake in terms of identifiable circuit motifs and generate testable predictions for targeting mPFC–insula–striatal– VTA interactions in alcohol use disorder.

## Introduction

Binge drinking refers to a high rate of alcohol consumption that results in blood alcohol levels of 0.08 grams of alcohol per deciliter in under 2 hours (*Understanding Binge Drinking | National Institute on Alcohol Abuse and Alcoholism (NIAAA)*, n.d.). Frequent binge drinkers exhibit increased levels of depression, stress, violent behavior, and impaired judgment when sober. They are also at an increased risk for alcohol use disorders and dependence (Addolorato et al., 2018; Hingson et al., 2005; Waszkiewicz et al., 2018). Significant progress has been made in identifying the brain mechanisms that control behavior within a binge session, which now makes a systems-level computational model feasible. The goal of the current study is to provide a computational model that integrates several of the currently identified brain mechanisms that govern binge drinking.

When initiating a binge, it is hypothesized that an individual vigorously seeks the rewarding properties of alcohol through a behavior referred to as front-loading (Ardinger et al., 2022). This drinking pattern is characterized by an initial, transient high rate of consumption, which is hypothesized to be driven by the acquired incentive motivational properties of alcohol. This leads to the hypothesis that neural circuits that control motivation and add vigor to behavior play a central role in this behavior. When front-loading is terminated, it is common for drinking to continue at a reduced rate, which has been conceptualized as “maintenance” drinking required to maintain blood alcohol at the desired level (Ardinger et al., 2022; Doyon et al., 2005). There is a substantial literature showing that, with extended training, alcohol seeking becomes progressively less sensitive to outcome devaluation and increasingly dependent on habit-promoting dorsal striatal circuits, particularly the dorsolateral striatum (Corbit et al., 2012, 2014). Therefore, while brain mechanisms that mediate the transition from front-loading to maintenance drinking remain unclear, we have conceptualized them herein as those that regulate the transition to the habitual control of behavior. In this model we explore the neural basis of these two stages of the binge - front-loading and maintenance.

### Circuitry involved in binge drinking

Here we review data to design a minimal circuit model that exhibits features necessary to facilitate binge drinking. The circuitry that contributes to binge drinking most likely extended beyond the circuits we include here. However, our goal is to provide an initial model of those most thoroughly characterized in binge-drinking and motivated behaviors, more generally.

### mPFC is involved in initiating/terminating enhanced seeking

Changes in the activity of the medial prefrontal cortex (mPFC) in response to drug/alcohol-paired cues has been well documented (Halladay et al., 2020; Myrick et al., 2004; Timme et al., 2022). In rodent self-administration paradigms, subsets of mPFC neurons increase firing prior to and shortly after instrumental actions such as lever pressing, indicating that mPFC activity participates in initiating reward-seeking behaviors and evaluating their outcomes (Burgos-Robles et al., 2013). In humans, functional imaging studies similarly show that mPFC activity correlates with alcohol craving and cue-induced motivation to drink (Myrick et al., 2004). Moreover, pathway-specific manipulations demonstrate that silencing mPFC projections to nucleus accumbens suppresses cue-induced reinstatement of alcohol seeking, providing causal evidence for its role in reward-directed behavior (Keistler et al., 2017). Therefore, the mPFC plays a key role in regulating reward seeking behaviors and, specifically, alcohol seeking behavior.

Importantly, the mPFC is not a unitary structure but contains heterogeneous neuronal populations encoding multiple task variables simultaneously, including action value, expected outcome, effort costs, and internal motivational states (Howland et al., 2022). Electrophysiological recordings reveal that distinct ensembles of prefrontal neurons represent cue identity, predicted reward, ongoing action selection, and feedback signals during goal-directed tasks (Hardung et al., 2017; Hillman & Bilkey, 2012). Specifically, the prefrontal cortex, particularly medial PFC, has been shown to track both the anticipated value of outcomes and the cost-benefit profile of continued effort (Clairis & Pessiglione, 2024). Multiple studies have demonstrated the mPFC’s role in considerations of effort, choice difficulty, and adaptive coding of response outcomes, relevant to flexible foraging behavior (Kolling et al., 2016; Patrick et al., 2021; Shenhav et al., 2013; Vassena et al., 2017). Accordingly, subjects with lesions to the dmPFC were specifically impaired at calibrating their level of persistence (van Geen et al., 2023). Reviews of prefrontal cortical circuits further emphasize that the mPFC integrates information about internal states and predicted outcomes to guide action selection and behavioral persistence (Goldstein & Volkow, 2011; Howland et al., 2022). This capacity to integrate internal state information with ongoing behavior makes the mPFC a plausible neural substrate for the internal “set point” signal proposed in our model.

During binge drinking, the brain alcohol levels lag behind the decision to transition from front loading to maintenance (Ardinger et al., 2022). Therefore, there must be an internal state estimate that generates an internal transition signal. There is increasing evidence that front-loading during alcohol drinking is not just a passive behavior but rather an effortful and motivated pattern of consumption (Ardinger et al., 2022). Therefore, we assume that during front-loading, agents engage prefrontal mechanisms to estimate the sufficiency of effortful drinking behavior to achieve a desired internal state. In the present framework, the SetPoint population represents a cortical estimate of cumulative effort and alcohol intake derived from striatal feedback, which is used to regulate the transition between high-vigor front-loading and lower-rate maintenance drinking.

Chronic alcohol exposure alters several aspects of prefrontal computation that are directly relevant to this regulatory function. Both human imaging studies and rodent experiments show that repeated alcohol use produces structural and functional changes in the mPFC, including altered synaptic plasticity, impaired inhibitory control, and disrupted representation of action– outcome contingencies (Goldstein & Volkow, 2011; Koob & Volkow, 2016; Linsenbardt & Boehm, 2015). Alcohol exposure can weaken prefrontal top-down control over striatal reward circuits and impair the ability of cortical networks to regulate reward-seeking behavior (Koob & Volkow, 2016; Timme et al., 2022). Such impairments are thought to contribute to the shift from controlled alcohol consumption to compulsive drinking observed in addiction models. Within the framework of our model, these changes may reduce the effectiveness of cortical SetPoint signals that normally terminate high-vigor alcohol seeking, thereby prolonging front-loading and increasing vulnerability to pathological binge drinking.

### Anterior insula is involved in sustaining behaviors, including drinking

Human neuroimaging studies identified sustained activity in anterior insula (aINS; referred to as insula below) across diverse tasks, consistent with a role in stable task maintenance rather than only transient cue detection (Dosenbach et al., 2006). More broadly, recent work argues that insula subregions are functionally differentiated, with posterior insula representing more primary bodily states and anterior insula integrating those signals with cognitive and affective information to guide behavior in the current or predicted situation (Centanni et al., 2021). This framework is particularly relevant for alcohol use, where anterior insula has been implicated in craving, alcohol cue reactivity, and loss of control over intake. In rodents, inhibition of anterior insula reduces alcohol drinking, including aversion-resistant or compulsion-like intake, while projection-specific studies show that anterior insula outputs to dorsolateral striatum contribute to binge-like drinking and front-loading-related circuit plasticity (Campbell et al., 2019; De Oliveira Sergio et al., 2021; Haggerty et al., 2022). Accordingly, in the present model we conceptualize anterior insula not as a generic “go” signal, but as a circuit element that stabilizes ongoing drinking behavior by sustaining action in accordance with internal state, motivational salience, and alcohol-related cue context.

### mPFC and Insula are interconnected

mPFC and aINS have been shown to co-activate during sustained behavior. Specifically, inactivation of mPFC neurons reduces neuronal activity in the insular cortex and also impairs initiation of persistent actions (Y. Wang & Sun, 2024). The rostral location of the prelimbic PFC that has substantial connections to the anterior gustatory insular cortex contains licking-modulated neurons (Horst & Laubach, 2013). Anterior insula stimulation activated mPFC as shown by fMRI and c-Fos immunohistochemistry (Haaranen et al., 2020). Finally, dense fibers in the opposite direction - from the insular cortex to the prefrontal cortex – has been documented as well (Mathiasen et al., 2023). Therefore, mPFC and Insular neurons are interconnected and co-activated during sustained behaviors, which may provide a mechanism facilitating the maintenance of seeking behaviors.

### Insular and mPFC projections are integrated in the striatum

Functional analysis studies have demonstrated significant projections from the insular cortex to the nucleus accumbens (NAc) (Ghaziri et al., 2018; Haaranen et al., 2020; Rogers-Carter et al., 2019; Wright & Groenewegen, 1996). Anterior insular projections to the NAc have been shown to drive aversion-resistant alcohol consumption in rats (Jaramillo, Agan, et al., 2018). mPFC projections to NAc are also implicated in alcohol drinking. Silencing of the mPFC → NAc pathway blocks cue-induced reinstatement of lever pressing in mice, thus, showing a decrease in alcohol seeking behavior (Keistler et al., 2017). The integration of information from the PFC by medium spiny neurons (MSN) of the NAc has been shown to be altered in binge drinking mice (Kolpakova et al., 2022). Glutamatergic projections to dorsal striatum are also implicated in alcohol drinking behavior (J. Wang et al., 2007). Specifically, anterior insula to DLS connections have been shown to regulate binge drinking and in particular front-loading (Haggerty et al., 2022). Therefore, the striatum integrates projections from the insula and mPFC to regulate alcohol consumption.

### Ventral striatal activity encodes vigor of responses

Ventral striatal activity, particularly within the nucleus accumbens (NAc), is linked to the vigor of motivated responding. Rather than functioning simply as a “reward center,” the NAc has been proposed to integrate cognitive and affective information to support action selection and to increase the efficiency and vigor of motivated behavior, especially when the appropriate action is uncertain or dynamically changing (Floresco, 2015).Consistent with this view, human instrumental-effort studies show that anticipatory NAcc activation tracks motivated effort expenditure, whereas dorsal striatal activation is more closely associated with lower-effort, habitual responding (Kroemer et al., 2014). In the alcohol domain, heightened NAcc activation during decisions involving greater potential reward and risk predicts earlier onset of binge drinking in adolescents (Morales et al., 2020). Alcohol exposure may also strengthen NAc output pathways that promote responding: following binge-like drinking, cholinergic regulation of glutamatergic transmission onto NAc D1-type medium spiny neurons is shifted toward excitation, selectively increasing glutamatergic drive onto this response-promoting population (Kolpakova et al., 2022). Together, these findings support the view that NAc activity contributes to high-vigor motivated actions, including risk- and reward-driven alcohol-related behavior.

### Dorsal striatum is implicated in habitual and history-dependent drinking

Dorsal striatal circuits, in contrast, are most consistently implicated in the acquisition and expression of habitual alcohol seeking and drinking. The dorsal striatum is central to action selection and habit formation, with medial and lateral subdivisions contributing differentially to goal-directed and habitual control, respectively (Corbit & Janak, 2016). In alcohol self-administration models, extended training promotes inflexible alcohol seeking, and dorsal striatal manipulations alter the balance between flexible and habitual alcohol-directed behavior (Corbit et al., 2012; Corbit & Janak, 2016). More recent studies indicate that dorsolateral striatal involvement in binge-like drinking and front-loading is drinking-history dependent: DLS AMPA receptor antagonism affects binge-like alcohol drinking early in exposure and gates alcohol front-loading after alcohol history, while anterior-insular inputs to the DLS contribute to the maintenance of binge drinking (Bauer et al., 2022; Haggerty et al., 2022). Thus, dorsal striatal activity should be framed primarily as a mechanism for habitual or history-dependent alcohol actions, whereas ventral striatal/NAc activity is more directly linked to high-vigor motivated responding.

### VTA signaling encodes reward prediction in alcohol seeking and consumption

Dopamine (DA) release from VTA DA neurons in response to a conditioned stimulus dynamically reflects the expected reward (Schultz, 1997) (Mohebi et al., 2019). Pharmacological or optogenetic silencing of VTA DA neurons reduces both alcohol self-administration and cue-induced reinstatement (Budygin & Weiner, 2015; Rodd et al., 2004). In alcohol self-administration, striatal DA concentration increases during the first five minutes of alcohol consumption and the peak positively correlates with the ethanol intake during the first ten minutes of the drinking period (Doyon et al., 2005). Time at which dopamine concentration peaked corresponds to the time of peak consumption, although blood alcohol levels (BAC) peaked more than 40 minutes later (Doyon et al., 2005). Therefore, VTA DA signaling respond to stimuli showing the availability of alcohol by peaking during the initial front-loading phase of consumption and retuning to baseline.

Integrating the above data, we design a model that replicates the front-loading and maintenance phases of binge drinking. The transition between the phases is obtained as internal switch in population-specific activity due to interactions among components of the modeled circuitry. By modulating parameters, we mimic alcohol use disorders (AUD). Mechanisms leading to AUDs revealed by this model may help motivate future research on the networks involved in binge drinking and find therapeutic targets relieving AUD conditions.

## Methods

### Model Structure

Our simulations are intended to model the processes that evolve in time over each brain region during a binge episode. By implementing the neural circuitry (Figure 1) and constraining the activity of each node to its known functions, as descried in the Introduction, our goal is to better understand the algorithm that is implemented across these circuits to control binge drinking.

**Figure 1:**
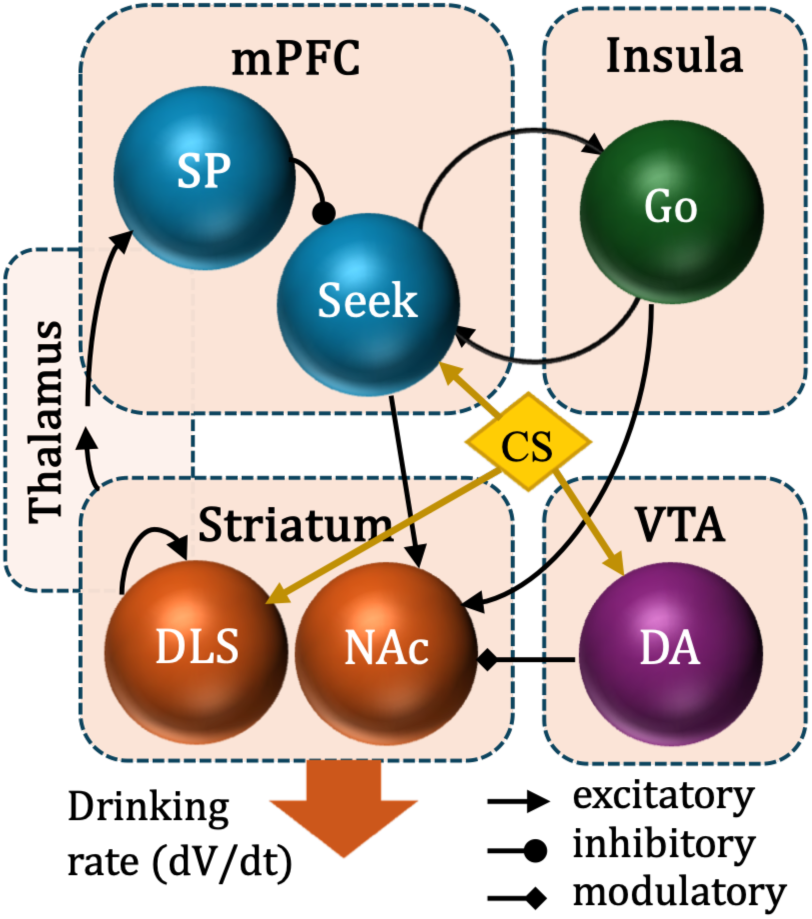
Schematic of the model circuitry. The following neural groups are included: SP=Setpoint, Seek=Alcohol seeking, Go=Persistent effort, DLS=dorsolateral striatum, NAc=nucleus accumbens, DA=VTA dopamine neurons. CS is the conditioned stimulus transiently applied at the start of a trial.

### Cortical Activity

The medial prefrontal cortex (mPFC) contains two neuronal populations. The first of them, Seek neurons, support alcohol seeking behavior and are initiated via a conditioned stimulus (CS). The Seek group is interconnected with an insular neural group involved in sustaining a behavior and called Go. These two groups mutually excite each other and therefore constitute a positive feedback loop. The second group in the mPFC called Setpoint (SP) reflects its estimation of the effort-benefit value. It is designed to integrate the alcohol consumption effort made during the trial, the information about which is originated from subcortical part of the model (see below).

### Subcortical Activity

The mPFC Seek and Insular Go neurons further project down to the nucleus accumbens (NAc), sending a sustained signal for the agent to maintain drinking behavior during the high-vigor front-loading phase of the binge.

In addition to the excitatory glutamatergic cortical projections, the NAc receives dopaminergic projections from the Ventral Tegmental Area (VTA). In alcohol self-administration, DA concentration in striatum increased during the first five minutes of ethanol availability, followed by a return to baseline (Doyon et al., 2005). Time at which dopamine concentration peaked corresponds to the time of peak consumption, although BAC peaked more than 40 minutes later. These results justify the transient spike in VTA activity in our model during the initial drinking period in response to the conditioned stimulus (CS). For simplicity, we model the CS as a sustained 3-minute session-onset alcohol-availability signal rather than as a brief phasic Pavlovian cue. This choice provided a gradual increase in DA for the duration of the CS. Accordingly, the CS duration should be interpreted as a computational approximation of sustained cue/context-driven dopamine accumulation during drinking initiation. The VTA activity is assumed to be proportional to the DA concentrations in the NAc, where DA modulates the firing characteristics of the neurons (Snyder et al., 2000).

Together with the NAc, we included a neural population reflecting dorsolateral striatum (DLS) activity. Its activation is important for maintaining habitual behavior and, therefore, provides a low vigor seek signal throughout the entirety of the drinking period. The DLS is activated by the CS in the model and remains active till the end of the simulation due to self-excitation that mimics its mutual excitatory connections with sensorimotor cortices and other structures (Aoki et al., 2019).

### Alcohol Consumption

The vigor of drinking is controlled by the striatal activity, where the NAc mostly contributes to high vigor episodes, and the DLS defines the low-level background consumption rate. Therefore, their sum defines the total drinking rate *dV*/*dt* in the model (Fig. 1).

### Dynamics of Setpoint

The same striatal activity is integrated by a group of cortical neurons called Setpoint (SP), which therefore closely follows the consumed alcohol volume, or the effort made during the trial. The effort estimate is assumed to be mediated by striatal-cortical connections in the model (Fig. 1) since basal ganglia-thalamo-cortical loop is an immediate candidate for such a role (Alexander et al., 1990). However, other connections that serve the function of relaying the information about the effort to the PFC cannot be excluded. Therefore, we omit specific details about this connection and made it purely functional. Thus, the Setpoint neural group tracks the agent’s overall effort and alcohol consumption and inhibits the activity of the Seek neuron group. The inhibition is nonlinear and surges once an internal threshold of the Setpoint activity is met.

### Trial structure

A trial starts with a conditioned stimulus (CS) that alerts the agent of the availability of alcohol. The CS is conceptualized as a general external cue/context associated with alcohol (not specific to any stimulus domain) and initiates alcohol seeking behavior (Ginsburg et al., 2022). The CS duration can vary across studies, but importantly the initiated behavior continues past its termination. The CS was directly applied as an input simultaneously to PFC Seek, DA, and DLS neural groups as these three groups are, among those important for initiating drinking, most likely to receive immediate stimulation from sensory cues (Doyon et al., 2005; Halladay et al., 2020). We ran our model over a simulated period of 120 minutes, consistent with ethanol access periods observed in front loading studies (Ardinger et al., 2022; Flores-Bonilla et al., 2021; Rhodes et al., 2005).

### Equations

We model the activities of neuronal populations with a continuous-time neural network. A well-established firing rate formalism (Cowan et al., 2016) is used to directly describe activity of each neural group by a differential equation. This formalism is most suitable for the task as firing rate is measured in most electrophysiological experiments as a proxy for the functional activation of neural groups. The time-dependent firing rate of neuronal population A is defined as follows (see Supplement for complete set of equations)

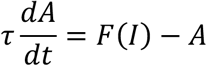

where *F* is an activation function, and *I* is a sum of the inputs from projecting neuronal populations (see below). τ is a time constant dictating the quickness at which the neuron group responds to a perturbation of inputs. The activation function F is sigmoidal and defined by:

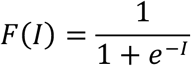

F normalizes firing rates on the unit interval [0,1] and introduces nonlinearity to the system.

The sum of inputs *I* accounts for the excitatory glutamatergic, inhibitory GABAergic projections among neuronal populations. For example,

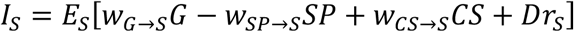

Where *I_S_* is the input to the Seek neurons, *E_S_* is the gain (excitability) of these neurons, *w_G_*_→*S*_ and *w_SP_*_→*S*_ are weights of projections from the Go and Setpoint groups, whereas *G* and *P* are their activation levels respectively. The *w_CS_*_→*S*_*CS* term describes activation by the CS, and the *Dr_S_* is the internal drive that defines baseline activity of this groups without inputs. The excitatory and inhibitory connections are taken into account by the signs of the corresponding input terms.

At each time step, a small normally distributed noise with variance σ = 0.3, *Z_i_* ∼ N(0, σ) is added to the equations. Applying Z on the input side of the activation function, i.e.

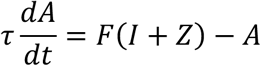

helps determine whether the dynamics of our model are stable with respect to small perturbations and produces more physiologically realistic firing characteristics.

The dopaminergic projections to the striatum are modulatory. They modulate the excitability, *E_N_*, and the drive, Dr*_N_* of the NAc neurons as follows (Snyder et al., 2000; Steephen, 2011):

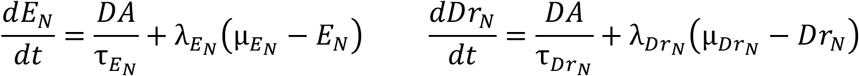

where λ_×_ represents the modulation and µ_×_ represents the baseline. According to these equations, these parameters revert to the mean once the DA concentration reverts to the baseline.

The main readout of the model, alcohol consumption (in g/kg) by the agent is represented by the integral of the striatal activity (NAc and DLS):

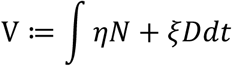

where *N*and *D* are NAc and DLS activity levels respectively, and *η* and *ξ* are scaling coefficients, which may be different for the two regions as they may contribute to the movement by somewhat different pathways.

### Model calibration

Current experimental techniques do not allow for direct measurement of parameters such as strength of connection between two neural groups. These values integrate multiple factors, including the number of neurons, input convergence, and synaptic current strength. This limitation is common in computational neuroscience, where models serve to integrate diverse data and generate testable predictions rather than precisely replicate biological circuits. The model parameters were selected through an iterative calibration process aimed at reproducing the following features; the two-phased drinking behavior, the internal switch between the phases, and the co-activation of mPFC and Insula beyond the CS duration. The parameter values were constrained by physiological plausibility and extensive data on the structure of the circuitry (see above). After successful calibration, the model was used to predict activity levels, circuit responses and functionality in conditions distinct from above experiments (e.g., modulation of projection strengths) and, in particular, reproducing physiological abnormalities. Furthermore, simulations were run to ensure robustness of the dynamics and present typical model behavior. While quantitative outputs may vary, the key patterns and trends remain consistent, indicating that the model’s conclusions are not sensitive to specific parameter choices.

### Analysis

In Results, we compute the activity of each neuron group by solving numerically the system of ODEs detailed in the previous section. The initial conditions and parameters are given in the Supplement. In particular, the Python `scipy` package solves the ODE system over 500 timesteps, corresponding to 120 minutes of trial time. To account for biological noise, we sample from a centered normal distribution and feed it into the sigmoidal activation functions of each neuron at each time step. The standard python packages such as `matplotlib` aid in the visualization of our results.

To detect transitions between dynamical phases in time series data, we employed a piecewise linear approximation–based change-point detection method. For each time series *y*(*t*), we assumed the presence of at most one transition characterized by a change in slope. The algorithm searches for a candidate change point *t_c_* by fitting two independent linear regressions to the data segments before and after it. For each candidate change point within a predefined admissible interval (excluding an initial transient period of 3 minutes and boundary regions), the sum of squared residuals (SSE) of the two linear fits was computed. The breakpoint minimizing the total SSE was selected as the optimal candidate.

To ensure that the detected breakpoint corresponds specifically to a reduction in slope, candidate change points were accepted only if the slope after the breakpoint was sufficiently smaller than the slope before the change point, exceeding a predefined threshold. If this criterion was not met, no change point was assigned for that time series.

In addition to the change point time, we quantified the magnitude of the transition by computing a *frontloading coefficient* (FL), defined as the ratio of the slope before the change point to the slope after the change point. This metric captures the relative change in accumulation rate across the detected transition.

To quantify the sensitivity of model outputs to individual parameter variations, we performed a one-at-a-time (OAT) sensitivity analysis. In this approach, each model parameter was perturbed independently while all other parameters were held fixed at their baseline values. For each parameter *p*, a set of multiplicative scaling factors was applied to generate perturbed parameter values of the form *p*′ = *f* ⋅ *p* , where *f* denotes the perturbation factor. For each perturbed parameter set, the model was simulated using the same initial conditions and time discretization as in the baseline case.

From each simulation, two summary metrics were extracted using the piecewise linear change-point detection procedure described above: (i) the transition time (frontloading duration) and (ii) the frontloading coefficient, defined as the ratio of slopes before and after the detected change point. These quantities were compared to their baseline counterparts to assess the effect of parameter perturbations.

Sensitivity was quantified by computing both absolute and relative changes in the detected transition time and frontloading coefficient with respect to the baseline simulation. Perturbations that did not yield a valid change point according to the slope-based acceptance criterion were excluded from sensitivity comparisons for the corresponding metric.

This OAT sensitivity analysis provides a local, interpretable measure of how individual parameters influence both the timing and magnitude of phase transitions in the model dynamics, while avoiding confounding interactions between parameters.

## Results

*The model reproduces binge drinking pattern* First, we reproduce behavioral data on the drinking pattern observed in binge drinking (Ardinger et al., 2022). Fig. 2 shows the volume of the consumed alcohol (in g/kg) over the trial time. The volume is approximated by the weighted integrated output of the striatum. The model successfully replicates the two-slope behavior of binge drinking commonly found in literature (Ardinger et al., 2022). Approximately the first 20 minutes is characterized by a rapid rate of drinking and constitutes the front-loading period. This phase is followed by the maintenance phase where the rate of consumption significantly decreases. The presence of these two phases, their duration and rates of consumption depend on the choice of model parameters (see below).

**Figure 2:**
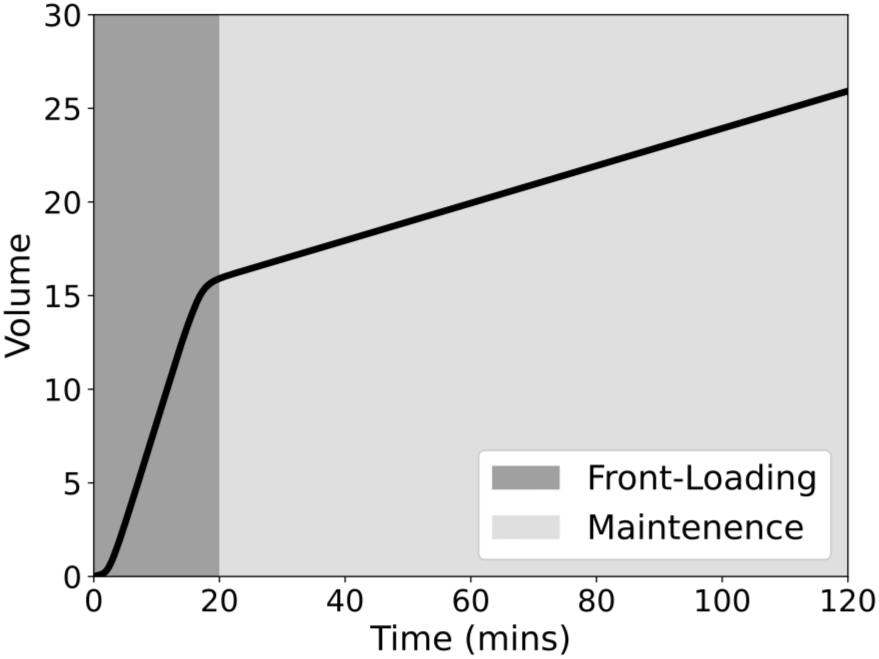
Drinking behavior is separated into two phases: high-vigor front loading and low-vigor maintenance.

### The PFC-Insular Loop Determines Front-loading Initiation

Fig. 3 shows dynamics of the activity of all neural groups in the model that underly the drinking behavior. In response to the CS starting the trial, the Seek neuron group in the mPFC rapidly increases its firing rate (Fig. 3A). The intracortical connections relay this activation to the Insular

**Figure 3:**
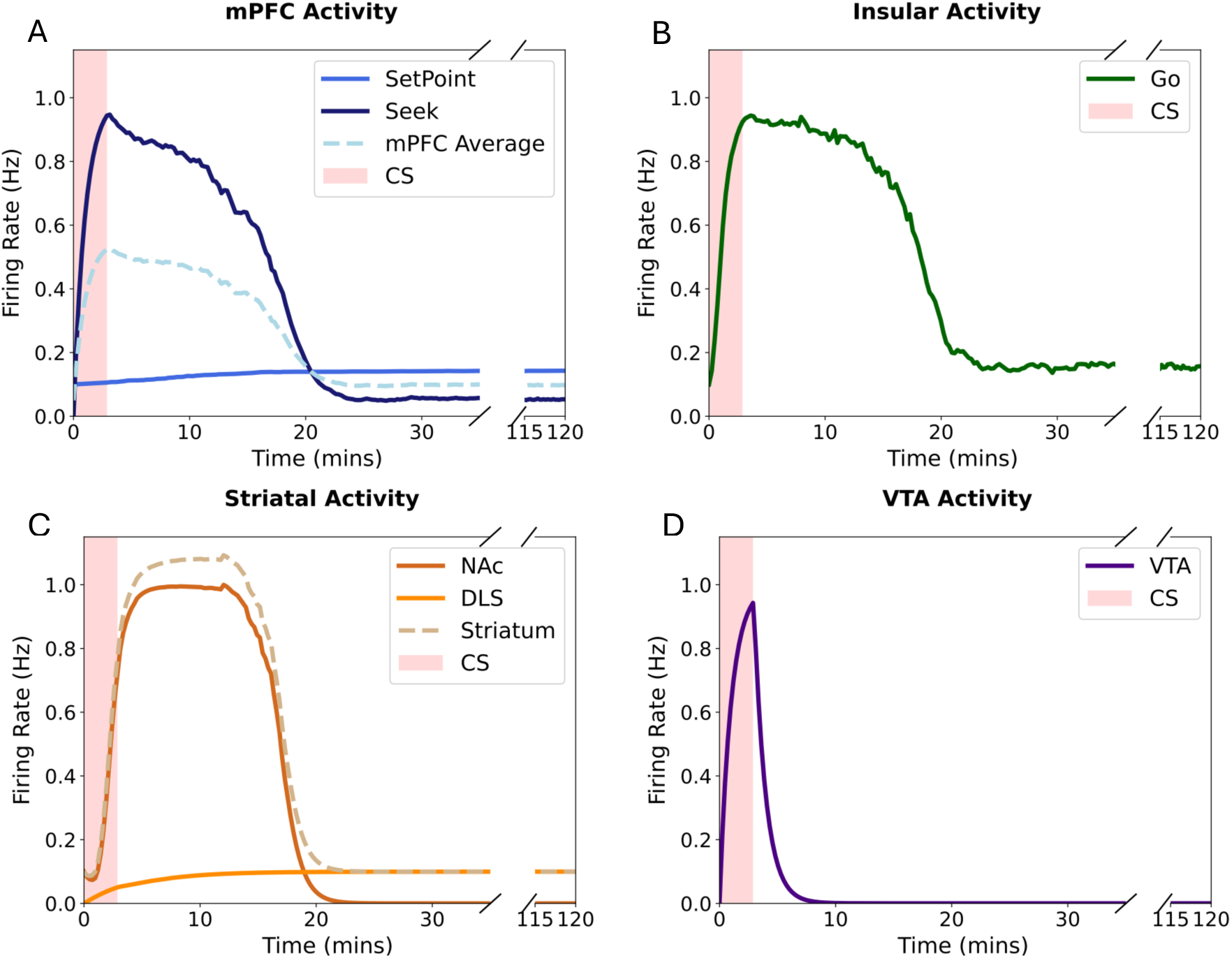
Activity patterns of all neural groups in the model during binge drinking. Notation and colors match the schematic in Fig. 1

Cortex, resulting in a rapid increase in firing of the Go neuron group (Fig. 3B). Due to the bistability established by the bidirectional excitatory connections between the mPFC and the insula, the two neuron groups retain a high level of activity during the initial phase of the binge past the duration of the CS. This activity excites the NAc (Fig. 3C) and causes front-loading phase of the binge (Fig. 2).

The bidirectional projections between the PFC and insula support the continuation of the agent’s current behavior. Thus, we find that the strengths of these projections determine the persistence at which the agent front-loads (Fig. 4). In agents whose PFC-insular loop is weakened, the front-loading period does not extend beyond the duration of the CS (Fig. 4, Unsustained). In agents whose cortico-insular loop is excessively potentiated, the front-loading phase extends infinitely (Fig 4, Persistent). For a medium interval of synaptic strengths, the desired transition from front-loading to maintenance is observed (Fig. 4, Two-phased). The transitions among these behaviors occur abruptly as the parameters reach a threshold value (Fig. 4 left.).

**Figure 4:**
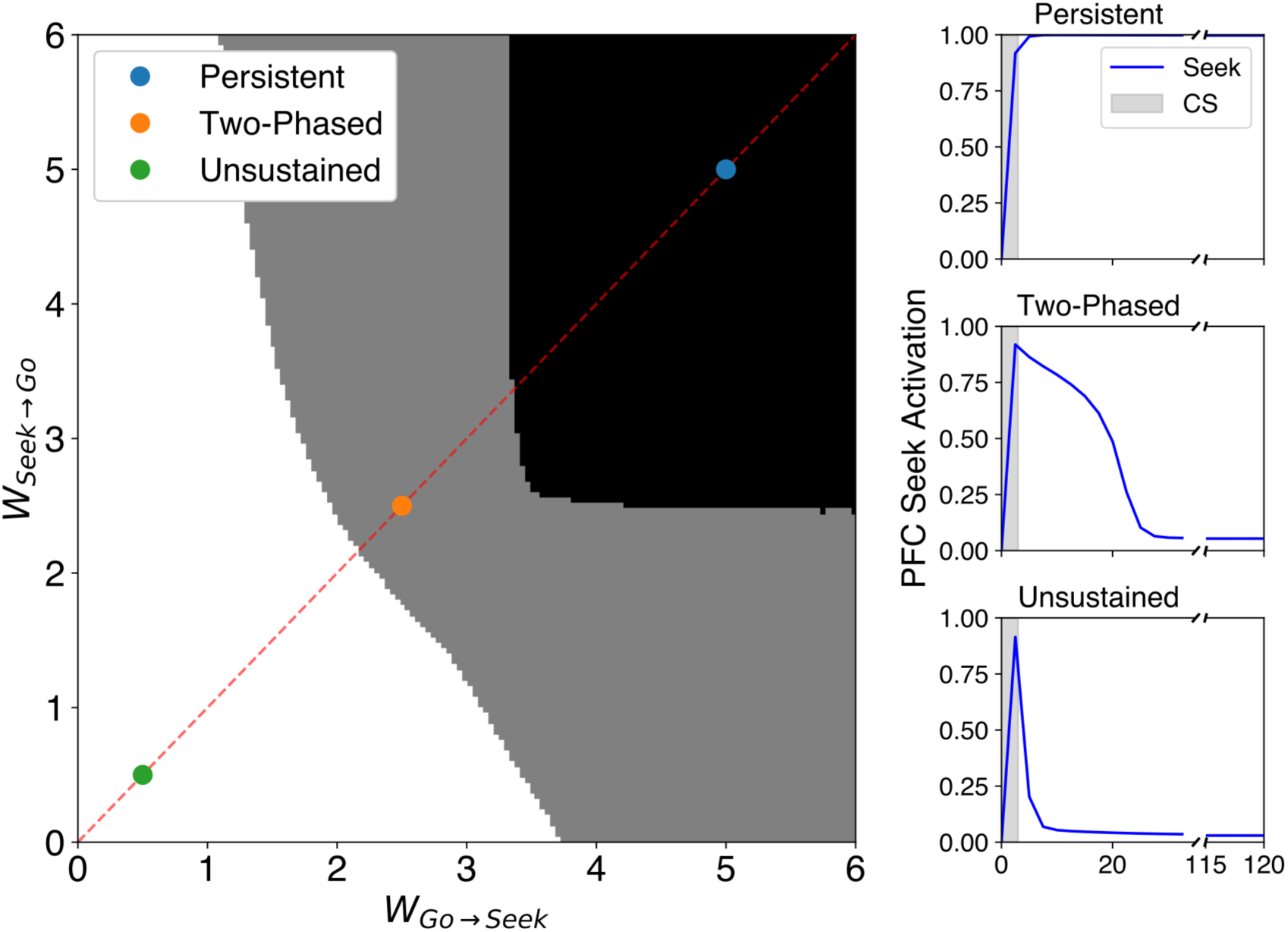
The connection between PFC Seek and Insular Go neural groups determine the dynamical regime of the whole circuit.

### Striato-Cortical Loop Determines Front-Loading Duration

The striatal activity representing alcohol consumption effort is integrated in the cortex by the Setpoint neural group (Fig. 2). We now examine what influence the strength of the feedback from the striatum to the cortex exerts on the behavior. Using change point analysis (see methods), we define the transition point from front-loading to maintenance phase (Fig. 2). This point defines the duration of frontloading. Fig. 5 shows that strengthening the projection from striatum to the Setpoint neurons in the cortex decrease the duration of the frontloading phase. Note that this modulation does not significantly change the intensity or vigor of front loading as the slope of alcohol accumulation remains the same (Fig. 5 inset). The strength of the other link in this feedback – the projection from the Setpoint to the Seek neurons in the cortex – exerts a very similar influence (data not shown). Therefore, the model predicts that the striato-cortical feedback determines the duration of the frontloading phase without affecting other aspects of the behavior such as its intensity.

**Figure 5:**
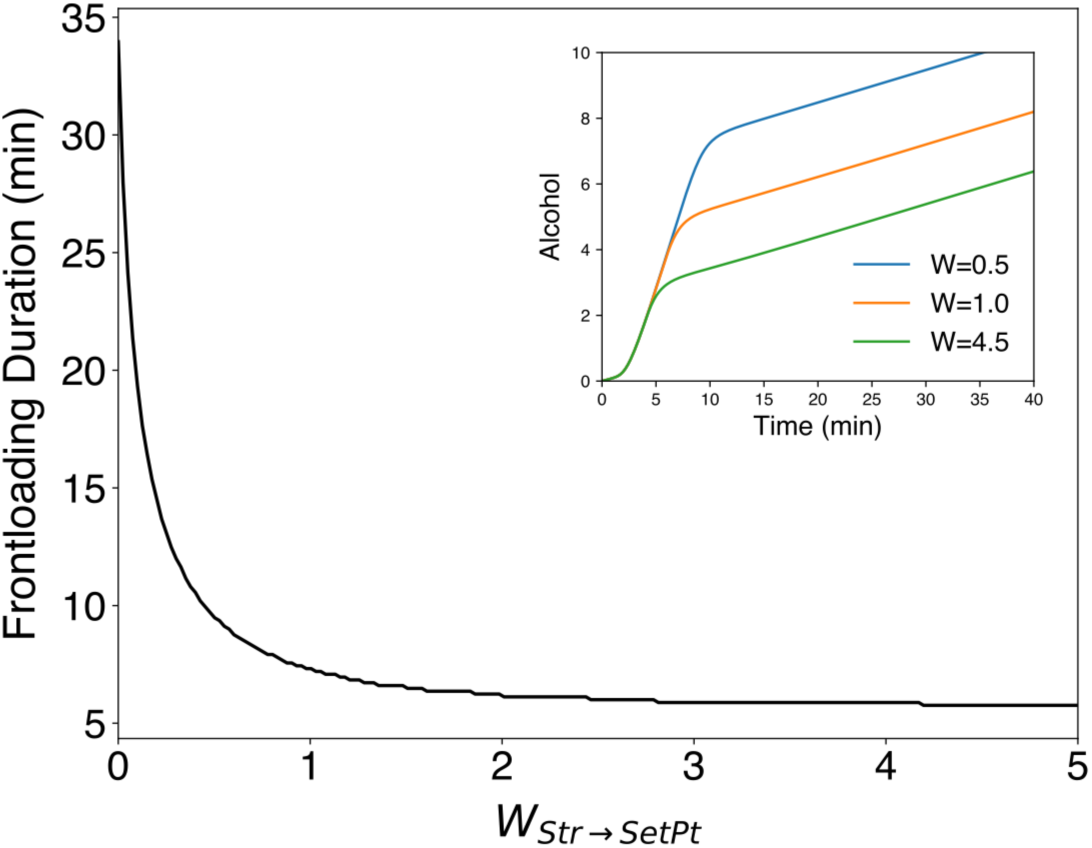
Duration of the frontloading phase depends in the strength of the striato-cortical feedback.

### Peak Dopamine Concentration determines the Intensity of Front-Loading

As reported in numerous studies, VTA activity increases during the first five minutes after introduction of alcohol (Doyon et al., 2005). In the model, VTA DA neuron firing peaks in response to the conditioned stimulus (Fig. 6 A). This modulation increases the excitability and drive of the NAc (see Model above). As a result, the NAc fires at an elevated rate during the initial phase of the trial (Fig. 6B). The elevated rate (Fig. 6C) is extended past the CS due to the slowness of the DA modulation processes in the NAc. Using the transition point *t_CP_* from front-loading to maintenance phase, we furher calculate the *Frontloading (FL) coefficient* as a ratio of alcohol consumption rates during frontloading and maintenance phases:

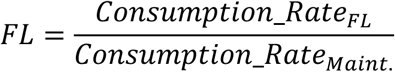

**Figure 6:**
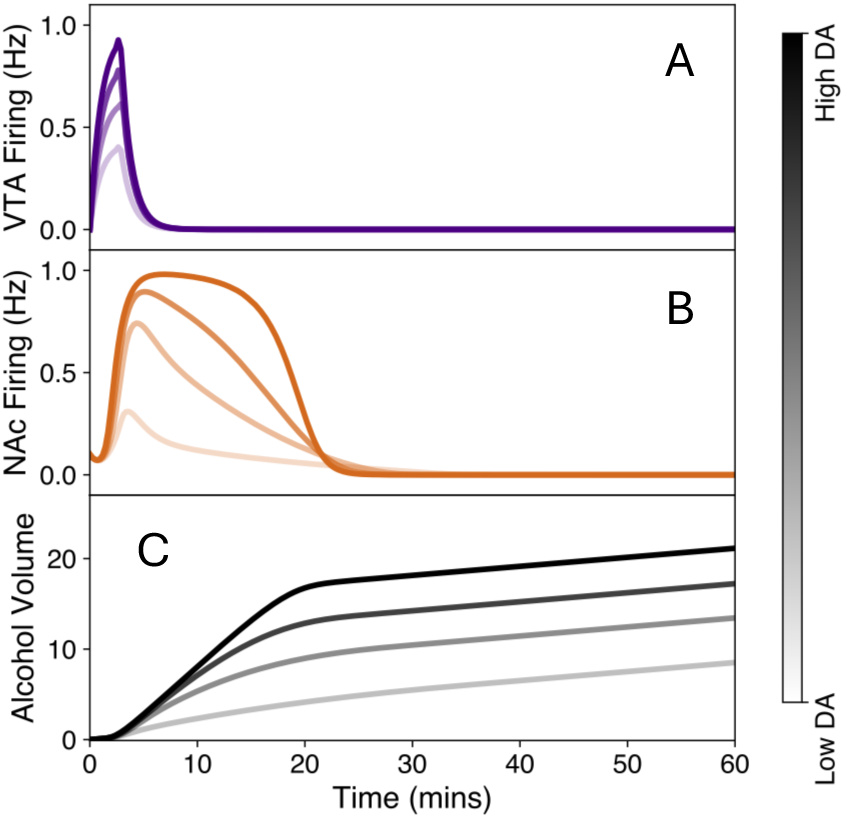
Front-loading intensity depends on the DA transient evoked by the alcohol CS.

Fig. 7 shows that the FL coefficient may increase several-fold as the total DA released by the CS amplifies.

**Figure 7:**
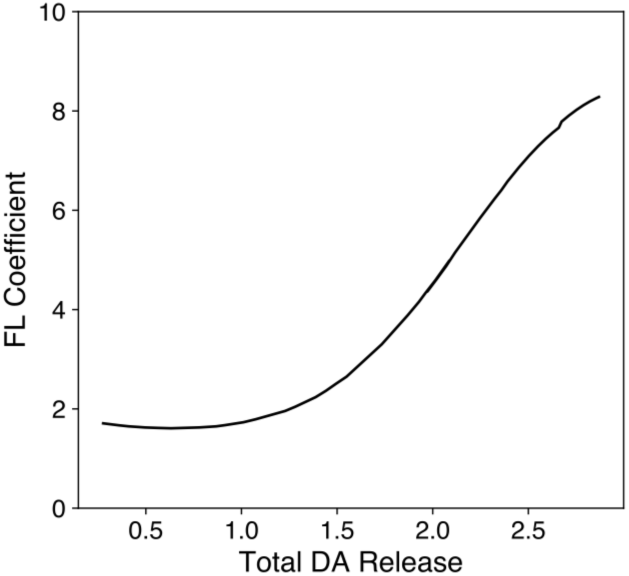
Frontloading coefficient increases as a function of the total DA released by the CS. The coefficient is computed as a ratio of alcohol consumption rates during frontloading and maintenance.

### Sensitivity Analysis groups modulations

As we show above, different features of the behavior such as the vigor and duration of the frontloading is differentially influenced by various components of the circuit. Next we undertake a systematic analysis of behavior sensitivity on the circuit parameters. Fig. 8 A and B show relative changes in the frontloading duration and intensity (the FL coefficient) respectively in response to modulation of synaptic weights among all neural groups in the circuit. The first groups of parameters consists of the connection weights between PFC Seek and Insular Go groups and the PFC SetPoint to Seek projection. All three weights affect both the duration and intensity of frontloading. Interestingly, if these parameters affect the duration monotonically (Fig. 8A), the FL coefficient decreases for most variations of the parameters (Fig. 8B). This is an indication of the transitions shown in Fig. 4: A decrease in these parameters virtually abolishes front loading, whereas their increase extends it infinitely, which abolishes maintenance phase and therefore reduces the FL coefficient without a significant reduction in the vigor of frontloading. Thus, these three paramters are the most influential because their modulation quickly compromizign the mechanism of the two-phased drinking pattern (Fig. 2).

**Figure 8:**
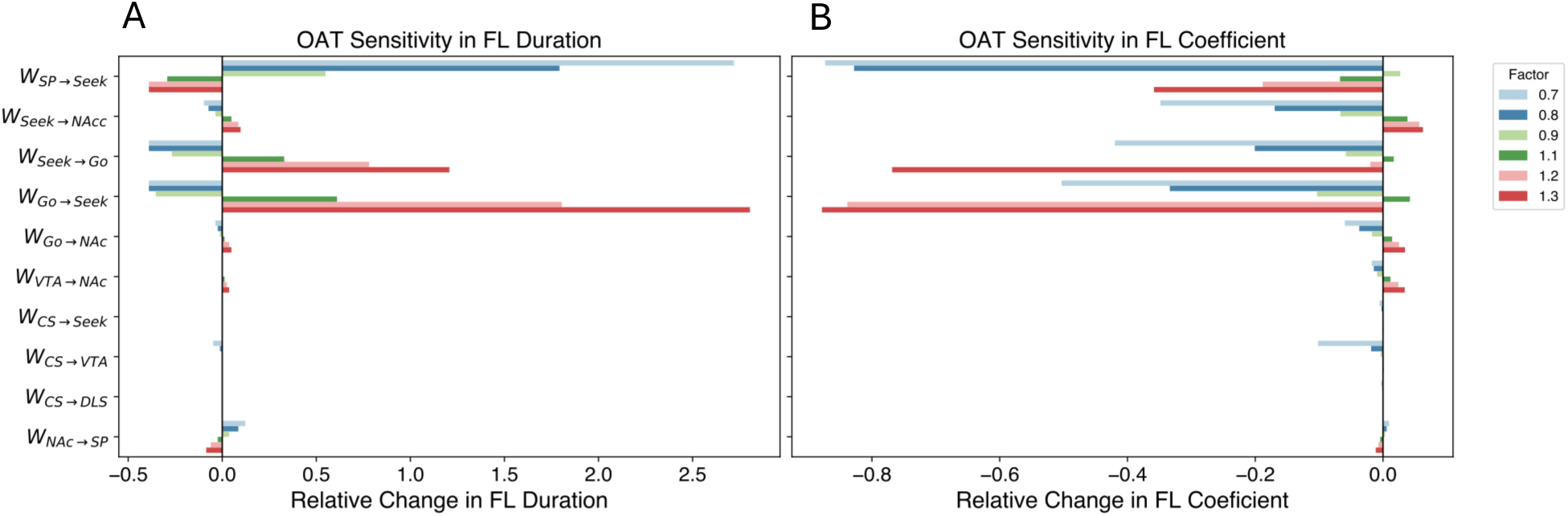
One-at-a-time (OAT) Sensitivity of the FL Duration (A) and FL coefficient (B) for each of the synaptic weights in the circuit.

The second group of paramters affect mostly the frontloading vigor (FL coefficient; Fig. 8B) rather than frontloading duration (Fig. 8A). These are weights of projections from both the Seek and Go groups to the NAc, VTA to NAc, as well as the strength of the VTA excitation by the CS. Obviously, these are all modulations that affect activation of the NAc. Whereas the Seek and Go groups directly excite the NAc neural group, the VTA DA pathway excerts modulatory influence on the activation.

The only paramter that affects the frontloading duration much more than its intensity is the weight of the feedback connection from the striatum to the PFC (SetPoint). This is because the excitation levels in the Seek-Go tandem are determined by the positive feedback loop between them, which creates bistability, rather than by an external signal. Such an external signal, when strong enough, flips the bistable switch and causes the abrupt transition from the frontloading to maintenance (Fig. 3).

## Discussion

The present model provides a mechanistic framework for understanding how interactions among the medial prefrontal cortex (mPFC), insula, striatum, and ventral tegmental area (VTA) give rise to the characteristic front-loading and maintenance phases of binge alcohol consumption. By integrating electrophysiological data and circuit-level findings into a dynamical systems model, we bridge behaviorally observed drinking microstructure with underlying neural computations.

### Integration of Model Predictions with Empirical Evidence

Our model identifies the **PFC–insular loop** as the key driver of front-loading behavior, consistent with evidence that these regions co-activate during motivated, sustained behaviors and alcohol seeking. Similarly, anterior insula activation enhances, and its inhibition reduces, voluntary alcohol consumption, particularly in aversion-resistant conditions (Campbell et al., 2019). The model’s demonstration that strong bidirectional PFC–insular excitation generates sustained high activity during early drinking mirrors this biology. Therefore, the model predicts the positive feedback loop between PFC and insula as promoting front loading pattern in alcohol drinking. In this framework, **front-loading emerges as a bistable regime**: a transient yet self-sustaining attractor state maintained by mutual excitation between mPFC and insula. This bistability explains why front-loading persists after the conditioned stimulus (CS) terminates, and predicts that perturbations such as cortical inhibition, attentional shifts, or interoceptive feedback can abruptly collapse it into the maintenance state. The ability of such perturbations to alter alcohol seeking and drinking has been shown before, but not for front-loading (Czachowski et al., 2003; Jaramillo, Van Voorhies, et al., 2018). The model predicts that cortical disinhibition or increased excitatory gain within the PFC–insular loop (e.g., through stress or prior alcohol exposure) would prolong or exaggerate front-loading, leading to pathological “persistent” states reminiscent of compulsive intake.

### Striatal Subcircuits and the Control of Vigor and Habit

The model dissociates **ventral and dorsal striatal roles** in binge drinking: nucleus accumbens (NAc) activity encodes vigor and motivational drive during front-loading, while dorsolateral striatum (DLS) contributes to habitual, low-effort consumption throughout the session. This distinction mirrors empirical data showing that NAc D1-MSNs display increased glutamatergic activation following binge alcohol exposure (Kolpakova et al., 2022), whereas DLS engagement strengthens with drinking history (Corbit & Janak, 2016). In our simulations, dopaminergic transients from the VTA drive the initial NAc activation, determining the intensity of front-loading, whereas striato-cortical feedback sets its duration. This aligns with reports that peak striatal dopamine release coincides with maximal drinking during early access (Doyon et al., 2005), suggesting that DA modulation temporally gates the high-vigor state. Moreover, our finding that stronger striato-cortical feedback shortens front-loading predicts that heightened feedback integration (e.g., via strengthened thalamo-cortical loops) could support adaptive termination of drinking, whereas weakened feedback may underlie prolonged binge episodes.

### Predicted Circuit-Level Transitions to Pathological States

The model’s parameter sensitivity analysis highlights three mechanisms by which physiological modulation could transition normal drinking into pathological binge states:

1. **Cortico-insular hyperconnectivity** causes persistent front-loading. Chronic alcohol use, stress exposure, or neuromodulatory changes that potentiate PFC– insular coupling may abolish the switch to maintenance. This maps onto imaging findings (Suk et al., 2021) showing hyperactivity and stronger functional connectivity within salience networks (including insula–PFC) in individuals with alcohol use disorder (AUD).
2. **Excessive VTA dopamine release** causes elevated front-loading intensity. Greater dopaminergic signaling amplifies the motivational value of cues, consistent with human fMRI studies where exaggerated ventral striatal activation predicts relapse risk and cue-reactivity in AUD (Heinz et al., 2004; Lim et al., 2020).
3. **Reduced striato-cortical integration** causes prolonged front-loading duration. In our model, weakened striatal feedback delays cortical inhibition of seeking, resulting in extended drinking bouts. This may reflect impaired prefrontal monitoring and diminished top-down control observed in chronic drinkers (Goldstein & Volkow, 2011).

Together, these mechanisms yield experimentally testable predictions:

- Chemogenetic dampening of insula→PFC or PFC→insula projections should shorten front-loading duration.
- Enhancing striato-cortical feedback (e.g., via targeted plasticity induction) should reduce transition timing.
- Manipulations that elevate phasic VTA DA (optogenetic or pharmacological) should increase early-session drinking vigor without altering its duration.

### Translational Implications and Future Directions

At the systems level, this work provides a computational scaffold linking **cue-driven motivation, cortical control, and striatal integration** to drinking microstructure. In human AUD, front-loading analogues manifest as rapid initial consumption and impaired ability to modulate intake. By mapping these behaviors onto network-level parameters—PFC–insula excitation, DA-mediated gain, and striato-cortical feedback—the model yields potential biomarkers and therapeutic targets. For example:

- Noninvasive neuroinhibition targeting anterior insula or dorsomedial PFC could attenuate pathological persistence by reducing loop gain.
- Pharmacotherapies that normalize dopamine modulation or enhance cortical feedback may moderate drinking by reducing subcortical and boosting cortical control.
- Neuroimaging studies could test model predictions by examining whether dynamic insula–PFC coupling and striatal BOLD amplitude predict real-time drinking rate in laboratory paradigms.

## Conclusion

By delineating how distinct circuit motifs govern vigor, persistence, and feedback control in binge alcohol intake, our model provides a unifying account of front-loading and maintenance behaviors. It generates explicit, experimentally testable predictions about how modulation of the PFC–insula–striatal network can either promote or suppress pathological drinking. Future work integrating sex differences, chronic exposure effects, and adaptive plasticity mechanisms will further refine this computational framework and enhance its translational value for understanding and treating alcohol use disorders.

